# A prosocial character of head-gaze aversion in common marmosets

**DOI:** 10.1101/2021.06.28.450151

**Authors:** S. Spadacenta, P.W. Dicke, P. Thier

**Affiliations:** Cognitive Neurology Laboratory, Hertie Institute for Clinical Brain Research, University of Tübingen

**Keywords:** common marmoset, eye contact, gaze aversion, social interaction, head-cocking

## Abstract

Gaze aversion is a behavior adopted by several mammalian and non-mammalian species in response to eye contact and usually interpreted as reaction to perceived threat. Unlike many other primates, common marmosets (*Callithrix jacchus*) are thought to have high tolerance for direct gaze, barely exhibiting gaze avoidance towards conspecifics and humans. Here we show that this does not hold for marmosets interacting with a familiar experimenter who suddenly establishes eye contact in a playful interaction (“peek-a-boo”). In video footage synchronously recorded from the two agents, we found that our monkeys consistently alternated between eye contact and head-gaze aversion. The striking similarity with the gaze aversion’s dynamics exhibited by human infants interacting with their caregivers suggests a shared behavioral strategy to disengage temporarily from overwhelming social stimulation, in order to prepare for a new round of rewarding, affiliative face-to-face interaction. The potential of our finding for a marmoset model of autism is discussed.

## INTRODUCTION

Establishing eye contact is a highly communicative act that shapes the social interactions of both humans and non-human primates [1]. Most primates perceive direct gaze as a display of threat preceding an attack [2,3,4,5,6,7,8], although eye contact, though brief, can also occur in prosocial contexts, such as in courtship, cooperative actions and play [9,10,11,12,13]. Eye contact, when sought and sustained, is more characteristic of mother-infant interactions across a wide variety of primate species, even the ones which, in adulthood, make use of direct gaze mostly in an agonistic context [14,15,16,17,18,19].

Irrespective of the different behavioral meanings that eye contact assumes according to species and contexts, across all human and non-human primates and many other mammalian species [20,21], after a varying period of direct gaze, a typical response to eye contact is to close the eyes or to turn the eyes or head away. This attempt to evade direct gaze is usually called “gaze aversion” [4,22]. It has been suggested that by means of this behavior (and also others aimed at covering the eyes), primates try to cut off the perception of arousing stimuli (e.g. direct gaze of a dominant animal) in order to continue ongoing activities, which would be impaired by excessive arousal [21]. A complementary possible function is that disengaging from eye contact may also signal appeasement, preventing an attack. In humans, gaze aversion is part of the normal behavioral repertoire of both adults and infants [23,24] and in line with the more flexible significance of mutual gaze shaped by context and interactor, it may as well assume different meanings. The role of gaze aversion as a regulator of perceptual input is particularly compelling in human infants [25], given their limitations to interact with the environment and to select or refuse visual stimulation. Cohn and Tronick [26,27] showed that infants’ gaze aversion is part of structured cycles of engagement (eye contact, smiles, etc.) and disengagement (gaze aversion, cry, etc.) when interacting with their caregivers. Human infants exhibit gaze aversion in reaction to the experience of the direct gaze of an emotionally unresponsive caregiver (“still face” experiment) [26,28], but also in playful interactions, for instance when playing peek-a-boo [29,30,31]. The notion that gaze aversion normalizes the infant’s level of arousal is suggested by the fact that, at least in the former case, looking away quickly normalizes elevated heart rate levels [29]. By the same token, gaze aversion may also serve as regulator of arousal due to positive affects [30]. To pause from eye contact, a source of emotional stimulation [32,33,34,35] seems to allow the infant to avoid a too distressful over-excitation and to recover for a new round of soothing emotional experiences provided by the caregiver’s face and eyes.

Common marmosets, a new world monkey species, are widely known to have a peculiar interest in faces [36,37] and to readily engage in mutual gaze in prosocial contexts, like for example when cooperating in joint actions [13]. Yet in common marmosets as well as in other new world monkey species, a use of gaze aversion in the regulation of social interactions, as exhibited in particular by human infants, has never been shown. Indeed, common marmosets are traditionally believed to barely make use of gaze aversion, arguably because they provide little indication that they may experience gaze as threatening when in contact with familiar individuals. Building on the serendipitous observation that marmosets engage in a “peek-a-boo” game with a human agent, we present evidence that this species deploys consistent gaze aversion behavior of a kind that we believe to have a prosocial character, namely as way to control overwhelming emotions which, if not bounded, would jeopardize the maintenance of the social interaction. Prosocial gaze aversion exhibited by marmosets suggests evolutionary continuity with a key role of gaze aversion in human behavior.

## MATERIALS AND METHODS

### Subjects

We tested 16 common marmosets (Callithrix jacchus; group 1: 3 females and 3 males, mean age: 3.8 ± 2.7 years; group 2: 6 females and 4 males, mean age: 5.5 ± 1.8 years), home caged at the Center for Integrative Science of the University of Tübingen. At the time of the study, the animals belonging to group 1 were involved in experiments independent of the observations of natural behavior in the facility addressed here, requiring behavioral training outside the facility, which is why they were in extensive daily contact with the experimenter. The animals belonging to group 2 knew the experimenter from her regular visits of the facility. All the subjects were tested in the presence of other marmosets in the facility, visually, but not acoustically, separated. All subjects had been born in captivity and they were kept in in the husbandry at approximately 26 °C, 40%– 60% relative humidity and a 12 h:12 h light-dark cycle. Access to water was always ad libitum, while food intake was controlled when the animals were subjected to the protocols required by the unrelated experiments.

### Experimental setting and procedure

We recorded common marmosets’ behavioral reaction when interacting with a familiar experimenter in a “peek-a-boo” game. The animals were tested in small transparent boxes (size 24 × 26 × 26 cm), permanently attached to the front part of each cage, accommodating free transition between compartments. Only the frontal and the right side of the box were fully transparent for visual access. Importantly, for group 1, the right side of the box allowed the view of the facility’s kitchen window, behind which the experimenter and the animal care takers showed up every day to access the marmosets’ facility. When animals expected to be moved to the setup for behavioral training or had heard that somebody had entered the kitchen, they usually went into the transport box, directing their attention towards the window. This configuration allowed us to serendipitously notice the head-gaze avoidance behavior, when approaching the window before opening the door to take one of the animals out for training.

The testing was performed under two different conditions: barrier or no barrier (see supplementary movie 1 and 2). In the barrier condition the experimenter was hiding behind the facility’s door, showing her face from the window at a random pace (Box - window distance: 220 cm for monkey Fin, 284 cm for monkey Han and 350 for monkey Flo, Fer, Mir and Ugh). In the no barrier condition the experimenter was standing closer to the animals, avoiding eye contact by looking down towards the floor and establishing eye contact by moving the head upwards (Box - experimenter distance: 100 cm). The interaction in both conditions started when the individual marmosets were calmly and spontaneously sitting in the transparent box waiting for the experimenter to engage in eye contact. In both conditions the procedure was repeated until the animal spontaneously moved back from the box to the inside part of the cage. Each time the animal moved out of the box back into the cage a session was considered ended and a new one started when the animal was back in the box. The number of trials per session varied according to how long the animal spontaneously interacted with the experimenter. The animals usually started to enter the box less frequently roughly 30 – 45 minutes after the onset of the recordings and therefore these never ran for more than 1 hour per day. Group 1 (n = 6) was tested by one familiar experimenter in both conditions (monkeys Fin, Flo and Fer a total of 200 repetitions per condition were collected; monkeys Ugh, Mir and Han 100 repetitions). The same experimenter tested the additional group of animals (group 2) only in the no barrier condition (n = 10), given that the facility’s structure did not allow the realization of the barrier condition for every cage position. As the stereotypical reaction was extremely consistent across animals we collected only between 20 and 40 trials in this second group. Additionally, we repeated measurements with a second experimenter who was familiar with 5 animals belonging to group 1, in both the barrier and no barrier conditions (40 repetitions per condition per monkey).

The marmosets’ behavior was video-taped using one camera facing the transparent box while a second camera, mounted on the same stand, faced the window (barrier condition) or the experimenter (no barrier condition), taking the animals’ perspective. Videos from both cameras were recorded synchronously at a frame rate of 30 Hz. The software IC Capture 2.4 was used to merge the two video files for later analysis using the OBS 23.02 software.

### Video analysis and variables scored

Individual video frames were extracted from the recordings and manually inspected and quantified using tools developed in MATLAB R2019a. For each trial we identified an “eye contact event” as the first frame in which the animal and the experimenter were in mutual eye contact. We then calculated the latency between this event and the start of the head-gaze aversion (gaze aversion latency), identified as the first frame in which the monkey’s head shifted away from the eye contact position in any direction. We documented also the following behavioral events: head-cocking after eye contact preceding the head-gaze aversion, whole body movement (rotation of the trunk together with the head relative to the longitudinal axis, thereby exposing the back of the animal towards the experimenter) during head-gaze aversion; eye blinks; vocalization produced at eye contact, soon after the head-gaze aversion or in between trials. Although we did not record sounds, the vocalization types were easily identified by the mouth movement and jotted by the experimenter at the end of each live session.

## RESULTS

### Common marmosets consistently respond to eye contact with head-gaze aversion

Both with the barrier present or not, and with both experimenters, at eye contact the animals reacted with stereotypical patterns of head-gaze aversion, summarized in figure 1. The head movement, of the order of 45° to 90° degrees relative to the trunk and executed in different directions in the fronto-parallel plane was coupled with a shift of the eye gaze axis away from the observer, as the videos did not indicate any significant counter rotation of the eyes relative to the head. After an interval of varying length eye contact was resumed. In the vast majority of trials, we observed the simple aversion pattern (fig. 1 panel a). A second pattern, observed in a smaller percentage of trials, was characterized by the addition of a vocalization that occurred either after having established eye contact with the experimenter (figure 1, panel b), soon after the aversion or in between periods of eye contact. In any case, the animals produced only contact calls (phees or twitters, for definition see [38]; fig 1, panel d). In a third pattern (fig. 1, panel c), the head-gaze aversion could be preceded by a head-cocking, a rotation of the cranium along the naso-occipital axis while maintaining a fixed eye axis direction, towards the experimenter’s eyes. Moreover, independent of the specific pattern, in a large percentage of trials an eye-blink preceded the head-gaze aversion by roughly 33-66 ms (1-2 video frames) or occurred concomitantly with it (group 1 pooled percentages reported in fig. 1, panel e; for individual values see supplementary table 1 and 2), reminiscent of head movement associated eye blinks in humans and non-human primates [39,40] and also in a few other species (e.g. peacocks, [41]). The fact that the eye blink preceded the head movement in most cases speaks against the possibility that the eyelid closure might be a protective, reflexive response to the head movement evoked draught, eventually stimulating the cornea [39]. Rather, closing the eyes might be a complimentary reaction, supporting the head turn in rapidly eliminating stimulation by the experimenter’s direct gaze. Moreover, we occasionally observed that many marmosets might blink not just once but several times when establishing eye contact with humans, before and throughout the head turn duration. If this form of blinking has also a communicative meaning remains an open question.

**Figure 1.**
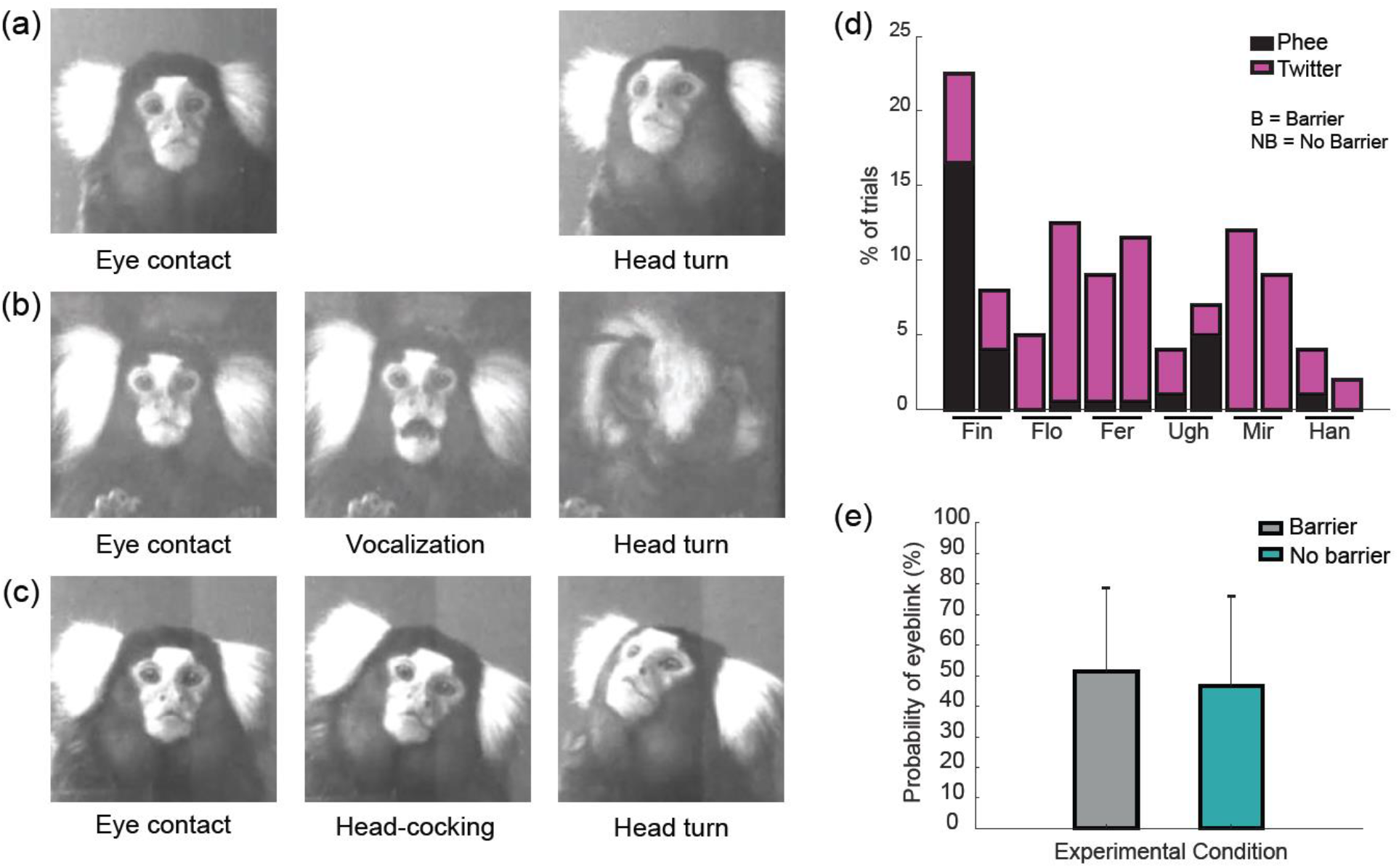
Patterns of response to eye contact and behavioral features. (a) Simple aversion. (b) A contact call was directed at the experimenter right after the eye contact. (c) Head-cocking (clock- or counter-clockwise) was exhibited after eye contact and before the aversion. (d) Percentage of trials in which a vocalization (phee or twitter) was produced. (e) Probability of eye-blink executed with head-gaze aversion (n = 6).

The head turns were usually carried out relatively smoothly and often slowly, or more rapidly, but very much unlike the typically fast and jerky gaze shifts serving the orientation to novel stimuli [42]. A presentation of the back in conjunction with the head movement was seen only occasionally and then primarily in paired animals, when the cage mate was simultaneously present in the transparent box (see supplementary table 1 and 2 under “body moved”).

We additionally analyzed head gaze aversion’s direction, considering 8 direction bins of 45° each in the fronto-parallel plane, as represented in fig. 2. In general, the preferred direction for aversion was the left side (animal’s perspective), namely towards the inner part of the cage. Moreover, 3 animals (Fin, Ugh, Mir) showed a shift of preference towards the down direction in the no barrier condition, where the animals had access to the experimenter’s gaze direction before eye contact. The result suggests that for these individual animals the final position of the head turn was influenced by the experimenter’s gaze direction, which we know from previous studies that common marmosets can follow geometrically [43].

**Figure 2.**
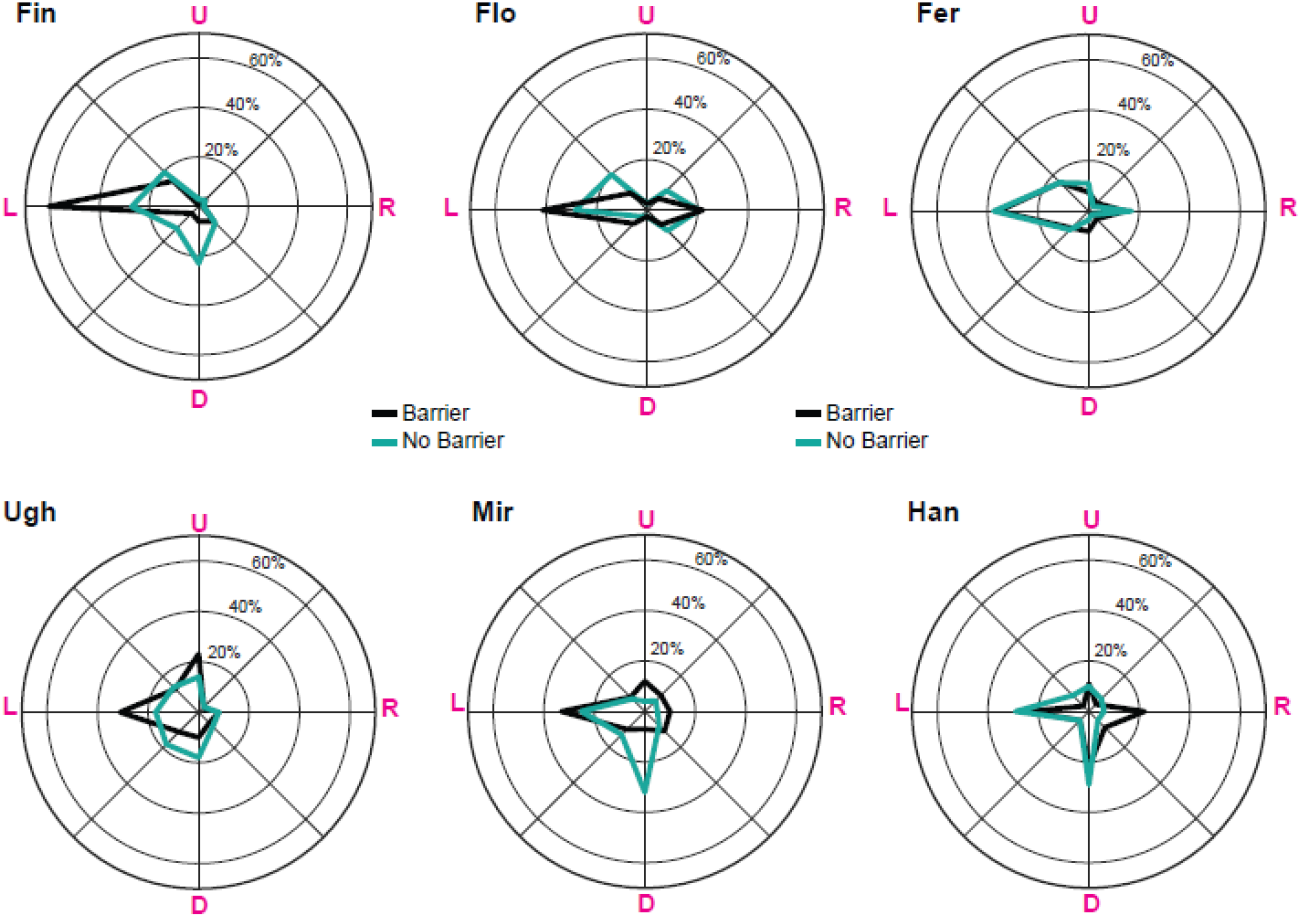
Head-gaze aversion directions (animals’ perspective) in the barrier and no barrier condition. L = Left, R = Right, U = Up, D = Down.

### Latency of aversion

The head-gaze aversion latency, or eye contact duration, defined as the time between the onset of eye contact and the initiation of the head movement, was in the large majority of trials below 1 second (see fig. 3 and supplementary table 1 and 2 for median values). For group 1, we explored the effect of the barrier presence on the aversion latency. Monkeys Fin, Flor, Fer and Mir averted significantly faster when the experimenter was in direct contact with them (Wilcoxon signed rank test, Fin: zval = 2.573, p < 0.01; Flo: zval = 5.874, p < 0.0001; Fer: zval = 4.926, p < 0.0001; Mir: zval = 4.829, p < 0.0001), monkey Han showed only a tendency in the same direction and monkey Ugh did not show any significant difference. Similar inter-individual differences of the barrier influence on the latencies were obtained when the second experimenter interacted with the animals (see supplementary fig. 1 and supplementary table 3). While the marmosets tested (Flo, Fer, Ugh, Mir and Han) exhibited exactly the same behavioral response patterns to the eye contact seen with experimenter 1, a barrier effect in the sense of a shortening of reaction times was seen in monkey Flo, Fer, Ugh and Han (although for the last one in the opposite direction) and absent in monkey Mir. Hence, the proximity of the experimenter in the no barrier conditions may shorten the eye contact duration but because of the profound interindividual differences, data for a larger group of animals would be needed to critically scrutinize this possibility.

**Figure 3.**
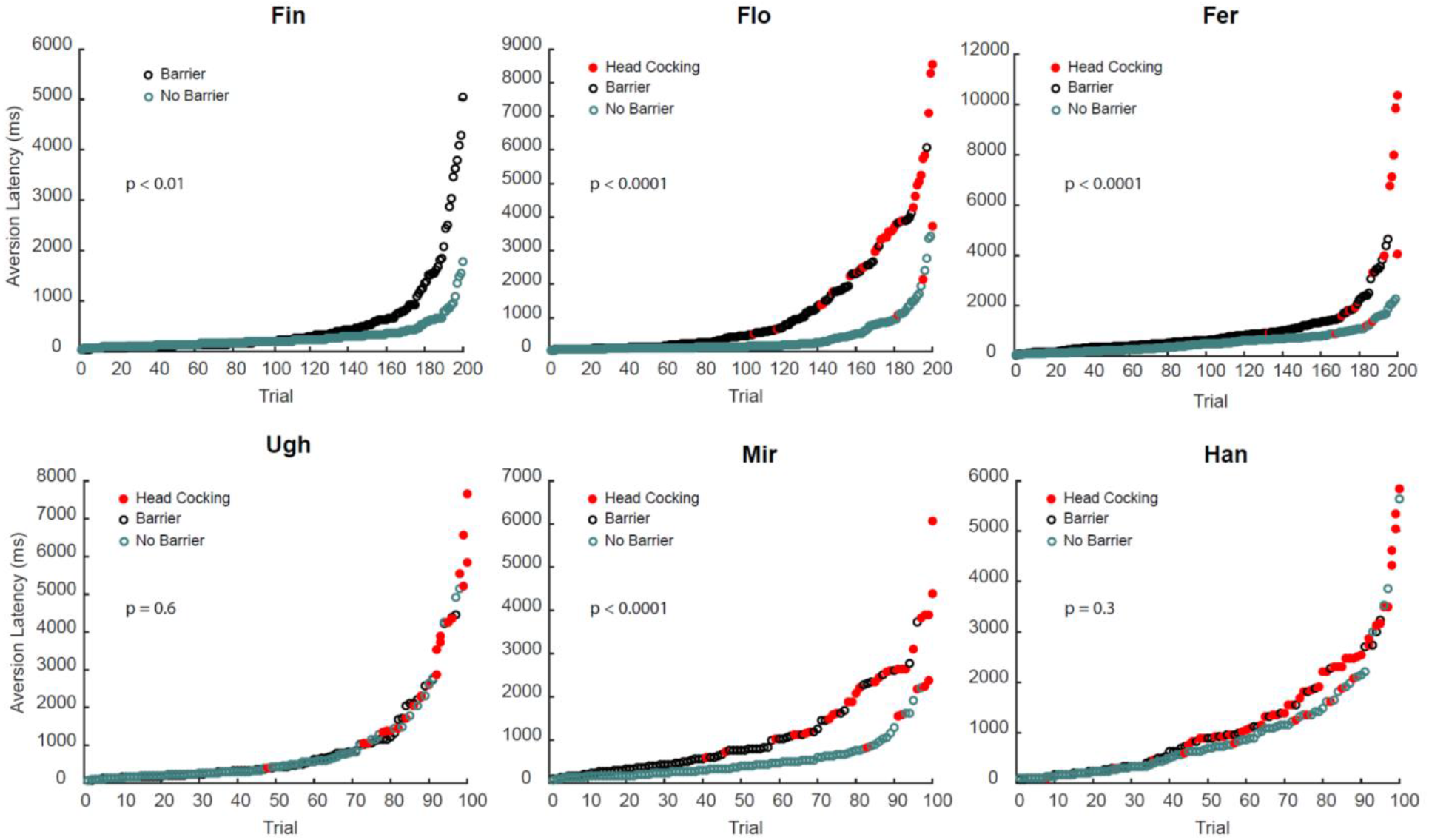
Effect of experimental condition on head-gaze aversion latency. For each monkey of group 1, we show the single trials aversion latency sorted by ascending duration. Red dots highlight trials in which the animals either performed a head-cocking or were looking at the experimenter with a tilted head position from the start of eye contact. The resulting statistics comparing barrier and non-barrier condition latencies with a Wilcoxon signed rank test are reported.

We then compared the aversion latencies between animals. In the barrier condition monkey Fin was significantly faster than all the others (Kruskal-Wallis test with Bonferroni correction, p < 0.0001), and monkey Flo significantly faster than Fer (p = 0.05) and Mir (p = 0.015). In the no barrier condition, still monkeys Fin and Flo turned out to be faster than all the others (p < 0.0001), but not significantly different from each other (p = 1).

Prompted by the observation that the first eye contact of each session, namely at the beginning of each play cycle, was longer as compared to the consecutive trial, we took a closer look at the dependency of gaze aversion latencies on trial number. When trial 1 was followed by a trial 2, the first eye contact duration was largely and consistently longer than the second one following the first head-gaze aversion, in both experimental conditions and for each individual animal, with only two exceptions in the no barrier condition (see figure 4; Wilcoxon signed rank test). For sequences of three or more trials, only monkey Flo showed a further shortening of the eye contact duration between trial 2 and 3.

**Figure 4.**
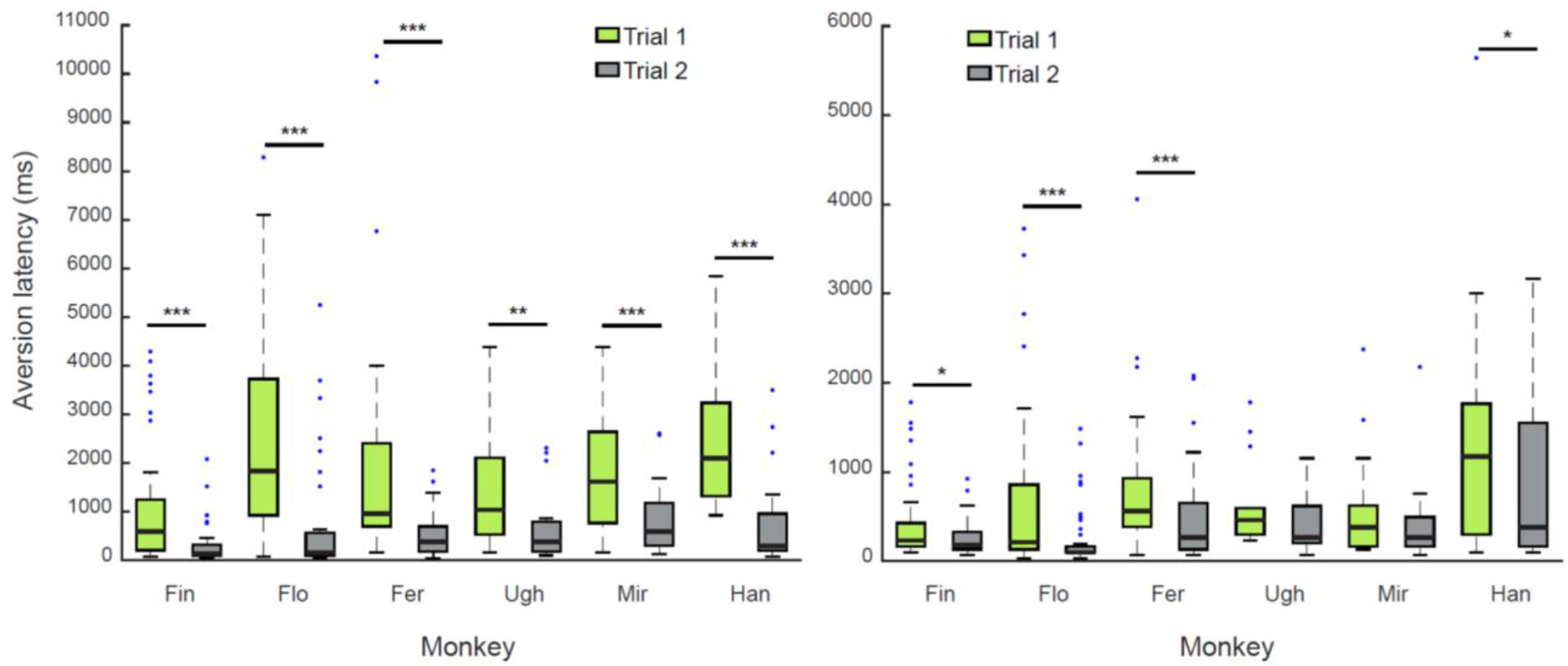
Comparison between head-gaze aversion latencies of trials 1 and trials 2. Eye contact duration on trials 1 was consistently longer than eye contact duration on trials 2. Only monkey Ugh and Mir showed no significant difference between type of trial in the no barrier condition. *p < 0.05, **p < 0.01, ***p < 0.001.

### Head-cocking influences head-gaze aversion latencies

As shown in fig. 1 (panel c), some animals, before turning the head away, responded to eye contact with a head-cocking (see supplementary movie 2 for examples), by definition a rotation of the observer’s head around a relatively fixed naso-occipital axis. The rotation was exhibited either clockwise or counterclockwise without significant difference. Previous reports of head-cocking in marmosets and other primates [44,45,46] described that this movement involves a fast saccade-like counter rotation of the head back to the upright orientation. However, the head-cocking we recorded preceded gaze aversion without an intermediate return to upright.

To understand whether tilting the head from the upright position had an influence on eye contact duration, we compared the aversion latencies of simple aversion trials, where the head was maintained in an upright position until the head turn, with trials in which the animals performed head-cocking or established eye contact already with the head deviating from the upright position. For group 1, this analysis was restricted to the barrier condition, in which the animals performed the larger number of head rotations. We found that when the animals performed head-cocking or engaged in eye contact already with a tilted head position, eye contact was maintained for a significantly longer duration as compared to the simple aversion trials (see fig. 5 for group 1 and supplementary fig. 2 for group 2; Wilcoxon sign rank test, Flo: zval = −4.937, p <0,0001; Fer: zval = −2.312, p < 0.05; Ugh: zval = −2.240, p < 0.05; Mir: zval = −4.280, p < 0.0001; Han: zval = −5.397, p < 0.0001). Moreover, the head-cocking followed the eye contact with a short latency (see supplementary table 1 and 2 for individual animals’ values) suggesting that the eye contact was the critical event triggering this behavioral response rather than other visual factors.

**Figure 5.**
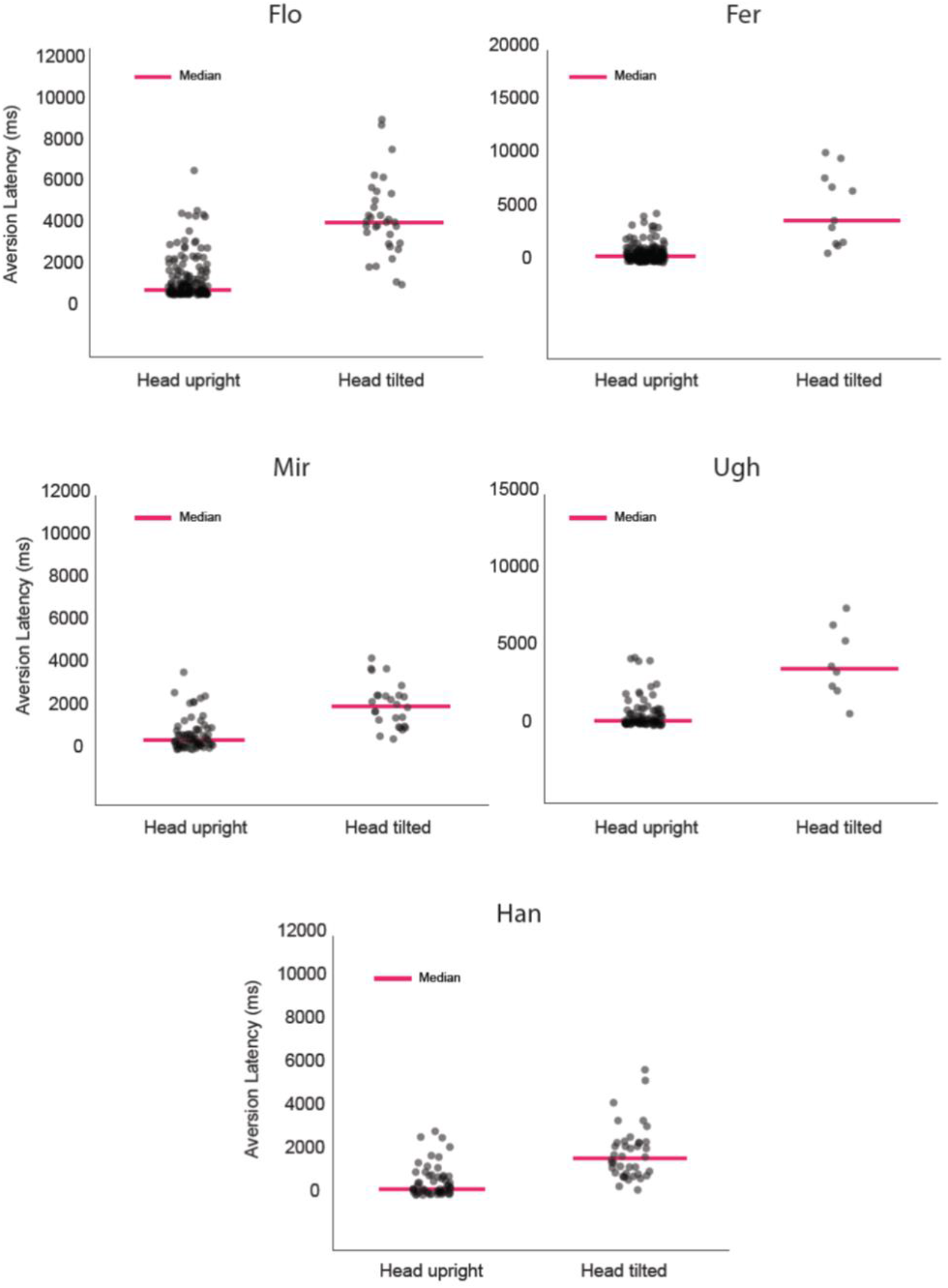
Maintaining eye contact with the head deviating from the upright position boosts eye contact duration. Single trial latencies of simple aversion trials (head maintained upright) and head-cocking trials (head tilted) are compared.

## DISCUSSION

We show that common marmosets consistently interrupt eye contact by means of a stereotyped head turn, when engaged in an interaction with a familiar experimenter who intermittently engages in eye contact with them (peek-a-boo game). Without doubt, looking at the faces of others, no matter if they are conspecifics or humans, and establishing eye contact is a rewarding urge for these animals, who have a strong preference for faces [37]. Yet, our observations demonstrate that direct gaze can only be tolerated for a limited amount of time, even in a familiar affiliative interaction, and needs to be temporarily interrupted by looking away. Given that direct gaze is rarely a form of threat for this species, also documented by the fact that our animals exhibited overall behavioral signs of positive arousal (contact calls, permanence in the testing box, absence of aggressive behaviors towards the observer), it is very unlikely that gaze aversion was an attempt to evade a felt threat or aggression like in many other primate species. Rather the urge to avert gaze must have a different reason. We suggest that common marmosets might break eye contact primarily driven by the need to cope with the emotional arousal elicited by direct gaze, while not perceived as threat or aggression still experienced as emotionally overwhelming. This is an interpretation that is congruent with the interpretation of gaze aversion as exhibited by human infants outlined in the introduction. Breaking eye contact time and again may help to bound the arousal level with the fundamental advantage of becoming able to prolong the overall duration of the pro-social interaction. The fact that marmosets exhibit a behavior strikingly similar to the one of human infants, still lacking executive control, might suggest that the disengagement is quasi-reflexively driven by subcortical structures, as a fast, protective mechanism against over-excitation.

We think that the type of emotional bond between animals and the familiar experimenter is the determinant of pro-social gaze aversion. The familiar experimenter is associated with positive experiences such as offers of food and treats as well as play. His/her direct gaze always signals a positive intent, which provides a strong motivation to interact, but also increases the level of arousal. A mechanism bounding the level of arousal will allow longer interactions, as the only escape from excessive arousal, a flight reaction, can be avoided. Arguably also the “peek-a-boo” behavior deployed by the experimenter to interact with the animals, in which visual access to her face and direct gaze was limited to distinct periods interrupted by pauses, has helped to prevent flight and maintain interest in the rewarding face and eyes. On the other hand, a flight reaction can typically be observed when common marmosets interact with an unfamiliar individual who stares at them for a prolonged time (intruder test). In this condition, experienced as threatening and dangerous rather than rewarding, an unbounded increase in arousal is certainly advantageous as it will elicit an early flight reaction, potentially vital to the animal’s survival. This reaction is accompanied by alarm calls, head and body bobs, and the avoidance of space closer to the other, spending longer time at the back of the cage [47,48], a behavioral pattern that is very different from the pattern of pro-social gaze aversion we observed.

The alternation of eye contact and gaze aversion will end after a few rounds rather than being maintained for longer. This may indicate that arousal levels can be bounded by gaze aversion only to some extent and may consequently accumulate over time, finally making it necessary for the monkey to stay away from the other. Indeed, the idea of incomplete arousal resetting is supported by our finding that the first eye contact in a given each session is always of longer duration as compared to subsequent trials. Of course, the alternation would also be ended at some point if the rewarding quality of the other’s face and eyes declined over time. Although a temporary decline of interest in the other cannot be excluded, the interest in the interaction must be rapidly restored as we did not observe any long-term changes in the attractivity of the human agent.

One may wonder if the head-gaze aversion that characterizes the interaction between a marmoset and a familiar human has relevance for interactions between marmosets in the absence of human interference. Indeed, we observed head-gaze aversion as elements of interactions of monkeys with conspecifics in two types of contexts. 1. During food competition: here, a subordinate animal, looking at a treat held by a dominant monkey, will avert its gaze as soon as the dominant animal engages in eye contact. 2. During re-union of familiar animals that had been separated for around 2 weeks. The latter configuration is reminiscent of the interaction with the familiar human and the behavior may be interpreted as an example of natural gaze aversion. The interpretation of the former configuration is less straight-forward. Here an attractor arguably inducing positive emotions – the treat – is around. On the other hand, the dominant monkey, in the possession of the treat will hardly be experienced as compliant to share and perhaps even as threatening. Hence, gaze aversion in this case is more similar to the standard agonistic patter exhibited by other nonhuman primates. Moreover, the subordinate monkey, by averting the gaze, would signal disinterest for the attractive treat, avoiding conflict.

### Head-cocking: a behavioral strategy to cope with eye contact?

The longest eye contact durations were registered when the animals looked at the experimenter, while assuming a head-cocking position before averting gaze. Does this observation suggest that head cocking may help to boost the tolerance to eye contact? Head cocking has previously been described as a stereotyped behavior exhibited by a large number of simian and prosimian primates [49] as well as by quite a few non-primate species (owls, dogs). Common marmosets are known to perform head-cocking in reaction to the appearance of new objects on the stage (e.g. flies, pieces of food), or other individuals (“head-cock staring”) like cage mates or human strangers [50]. It is more frequent when directed towards living objects [46] and it gradually decreases in frequency with age [45]. The functional significance of head-cocking in primates remains unclear. Time and again it has been suggested to facilitate the scrutiny of objects, in particular novel ones, by improving visual capacities [45,49,50]. Yet, the visual mechanism that might underpin this presumed role of head cocking in object analysis has remained elusive and experimental evidence supporting a role in visual perception has to the best of our knowledge never been presented. In our analysis, it emerged that head-cocking significantly prolonged the duration of eye contact with the experimenter. Hence, could it be that head-cocking helps to decrease the emotional impact of the other’s face and eyes, thereby allowing longer periods of direct gaze? We and other primates may quickly detect horizontally oriented eyes because of experience-dependent tuning of the visual system. By rotating the observer’s head, the retinal image of the pair will be tilted relative to the horizontal, arguably compromising perception of the *Gestalt* and consequently reducing its emotional impact. The fact that both humans and lemurs exhibit longer fixation duration when being exposed to tilted faces or tilted dots resembling eyes (humans: [22, 51]; lemurs: [4]) is clearly in line with the assumption that image tilt may reduce the impact of the other’s eyes. However, head-cocking may not only serve the animals’ arousal regulation, but may also help to stabilize communication by heralding a later, less abrupt disengagement based on head-gaze aversion. In any case, we emphasize that unfamiliarity with the human agent responsible for triggering head-cocking as suggested by previous work [52] can be excluded, as in our study the animals were used to see the same agent day after day, without exhibiting a decrease in head-cocking frequency over time.

## CONCLUSIONS

We have tried to provide evidence for a role of gaze aversion in ensuring bounds to the emotional impact of rewarding social stimulation offered by the face and the eyes of a familiar agent. The same function could be deployed by head-cocking directed towards other individuals. As the use of head-gaze aversion exhibited by a nonhuman primate whose evolutionary path started to deviate from us humans around 35 million years ago [53] is strikingly similar to its use by human infants engaged in dyadic interactions, the possibility of a shared and eventually even homologous behavior arises.

Alterations in the ability to engage in eye contact with a tendency to evade the other’s gaze are a hallmark of autism spectrum disorders (ASD), detectable already in early childhood during social games [54,55]. Our demonstration that marmosets and humans share a behavioral pattern compromised in ASD nourishes the hope that this monkey species may serve as useful model in future work on the roots of compromised eye contact in ASD.

## Ethics declarations

All the procedures were approved by the responsible national authority, the Regierungspräsidium Tübingen, Germany.

## Data availability

The datasets generated during and/or analyzed during the current study are available from the corresponding author on reasonable request.

## Author information

Affiliations

Cognitive Neurology Laboratory, Hertie Institute for Clinical Brain Research, University of Tübingen

## Contributions

S.S. conceived the study, performed the experiments and analyzed the data. S.S. and P.W.D built the experimental configuration. S.S. and P.T. interpreted the results and wrote the manuscript with contributions from P.W.D. All authors reviewed and approved the manuscript prior to submission.

## Acknowledgements

We are grateful to Elena Cavani for her help with data collection.

