## Supplementary information for "A prosocial character of head-gaze aversion in common marmosets"

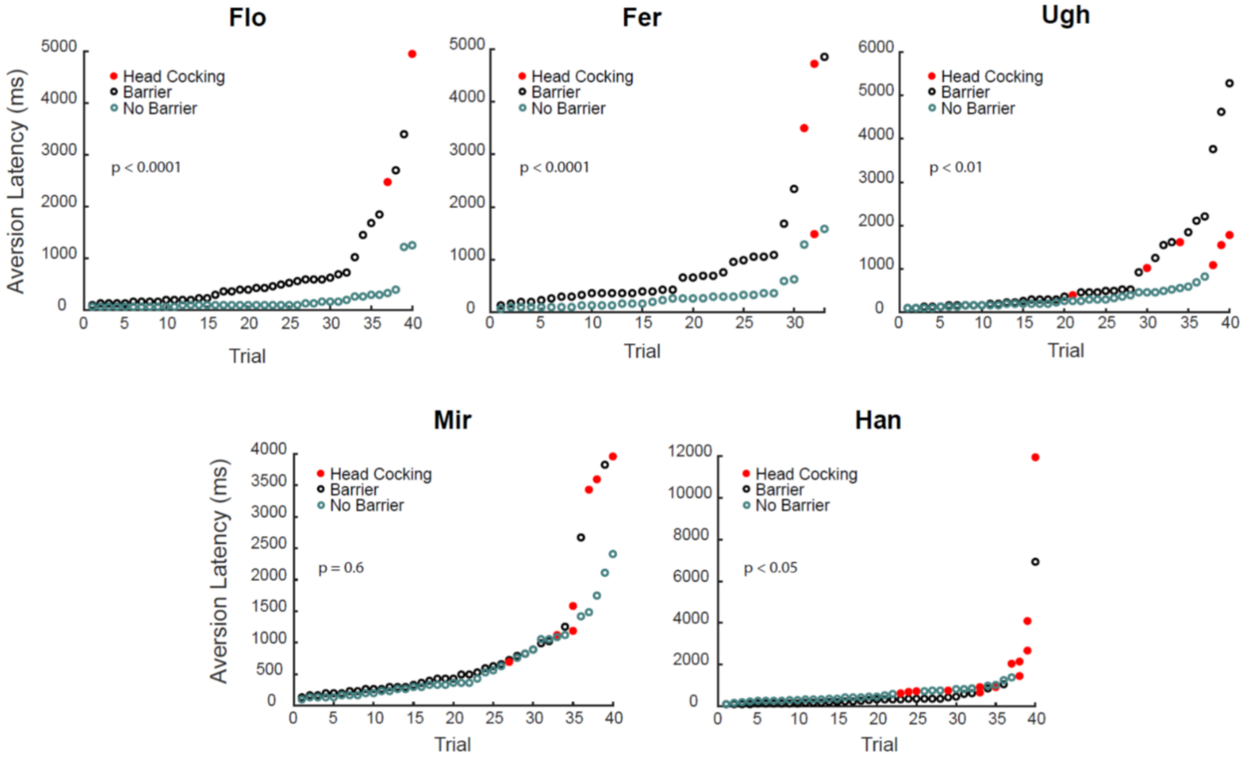


**Supplementary fig. 1**. Effect of experimental condition on aversion latency for 5 animals (Flo, Fer, Ugh, Mir, Han) interacting with a second familiar experimenter. For each monkey we show the single trial aversion latency sorted by ascending duration. Red dots highlight trials in which the animals either performed a head-cocking or were looking at the experimenter with a tilted head position from the start of eye contact. The resulting statistics comparing barrier and non-barrier condition latencies with a Wilcoxon signed rank test are reported.

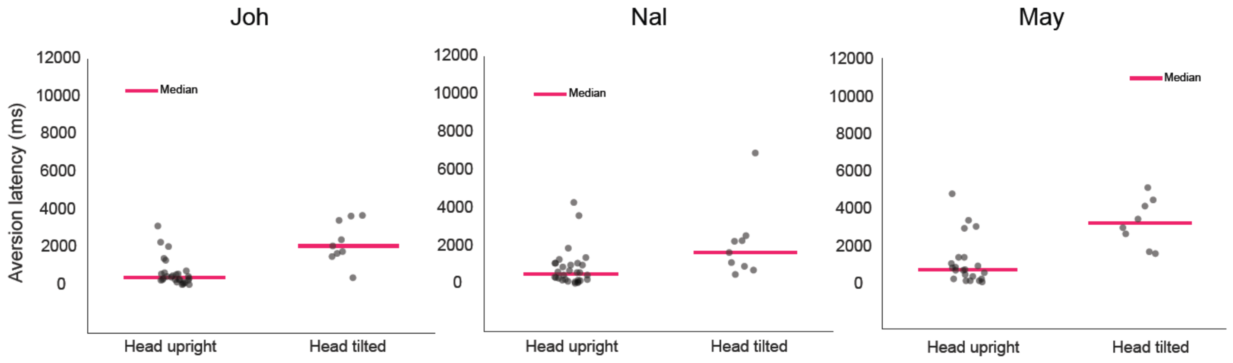


**Supplementary fig. 2**. Comparison between aversion latencies of simple aversion trials and head-cocking trials (Monkey Joh, Nal, May of group 2). As for group 1, eye contact was maintained for a longer duration when the animals head position was tilted (Wilcoxon signed rank test, Joh, zval = -2.521, $p = 0.01$; Nal: zval = -2.073, $p < 0.03$; May: zval = -1.96, $p < 0.05$). Each dot corresponds to a single trial latency.

| Monkey | Fin | | Flo | | Fer | | Ugh | | Mir | | Han | |
|---|---|---|---|---|---|---|---|---|---|---|---|---|
| **Condition** | Barrier | No Barrier | Barrier | No Barrier | Barrier | No Barrier | Barrier | No Barrier | Barrier | No Barrier | Barrier | No Barrier |
| Median aversion latency (ms) | 198 | 198 | 462 | 132 | 627 | 495 | 428.5 | 429 | 759 | 396 | 891 | 693 |
| Blinks (%) | 81.5 | 69 | 69.5 | 86.5 | 75 | 60.5 | 35 | 25.5 | 14 | 13.5 | 33 | 24.5 |
| Vocalizations (%) | 22.5 | 8 | 5 | 12.5 | 9 | 11.5 | 4 | 7 | 12 | 9 | 4 | 2 |
| Body moved (%) | 0 | 0 | 15 | 25 | 9 | 20.5 | 1 | 1 | 1 | 4 | 14 | 12 |
| Head- cocking (%) | 0 | 0 | 16 | 1.5 | 5.5 | 2 | 3 | 3 | 25 | 2 | 24 | 8 |
| Median head-cocking start latency (ms) | - | - | 198 | 1947 | 990 | 676.5 | 957 | 462 | 594 | 1666 | 429 | 907.5 |

**Supplementary table 1**. Behavioral values summary of group 1 monkeys.

| Monkey | Ali | Joh | Nal | Sis | May | Dai | Que | Eva | Wil | Giz |
|---|---|---|---|---|---|---|---|---|---|---|
| **Condition** | No Barrier | No Barrier | No Barrier | No Barrier | No Barrier | No Barrier | No Barrier | No Barrier | No Barrier | No Barrier |
| Median aversion latency (ms) | 198 | 528 | 726 | 297 | 957 | 1023 | 462 | 132 | 577.5 | 198 |
| Blinks (%) | 84 | 78,4 | 69 | 54.3 | 67 | 27.01 | 37.1 | 90 | 54.2 | 69.6 |
| Vocalizations (%) | 13 | 0 | 7.7 | 8.6 | 20 | 2.8 | 20 | 0 | 4.2 | 0 |
| Body moved (%) | 13 | 16.2 | 3 | 0 | 0 | 0 | 0 | 10 | 17 | 26.1 |
| Head- cocking (%) | 0 | 24.3 | 23.1 | 8.6 | 27 | 16.2 | 2.8 | 0 | 16.6 | 4.34 |
| Median head-cocking start latency (ms) | - | 528 | 429 | 561 | 511 | 297 | 495 | - | 165 | 495 |

**Supplementary table 2**. Behavioral values summary of group 2 monkeys.

| Monkey | Flo | | Fer | | Ugh | | Mir | | Han | |
|---|---|---|---|---|---|---|---|---|---|---|
| **Condition** | Barrier | No Barrier | Barrier | No Barrier | Barrier | No Barrier | Barrier | No Barrier | Barrier | No Barrier |
| Median aversion latency (ms) | 412.5 | 99 | 429 | 231 | 379.5 | 264 | 462 | 363 | 330 | 495 |
| Blinks (%) | 75 | 82.5 | 82 | 61 | 70 | 72.5 | 47.5 | 27.5 | 72.5 | 50 |
| Vocalizations (%) | 2.5 | 0 | 6.1 | 9.1 | 2.5 | 2.5 | 10 | 2.5 | 2.5 | 0 |
| Body moved (%) | 7.5 | 52.5 | 6.1 | 24.4 | 0 | 2.5 | 15 | 5 | 2.5 | 5 |
| Head- cocking (%) | 5 | 0 | 3 | 0 | 0 | 5 | 10 | 0 | 10 | 5 |
| Median head-cocking start latency (ms) | 891 | - | - | - | - | 181.5 | 1551 | - | 462 | 379.5 |

**Supplementary table 3**. Behavioral values summary of group 1 monkeys tested by a second experimenter.
